# Longevity defined as top 10% survivors is transmitted as a quantitative genetic trait: results from large three-generation datasets

**DOI:** 10.1101/373274

**Authors:** Niels van den Berg, Mar Rodríguez-Girondo, Ingrid K. van Dijk, Rick J. Mourits, Kees Mandemakers, Angelique A.P.O. Janssens, Marian Beekman, Ken Robert Smith, P. Eline Slagboom

## Abstract

Survival to extreme ages clusters within families. However, identifying genetic loci conferring longevity and low morbidity in such longevous families is challenging. There is debate concerning the survival percentile that best isolates the genetic component in longevity. Here, we use three-generational mortality data from two large datasets, UPDB (US) and LINKS (Netherlands). We studied 21,046 unselected families containing index persons, their parents, siblings, spouses, and children, comprising 321,687 individuals. Our analyses provide strong evidence that longevity is transmitted as a quantitative genetic trait among survivors up to the top 10% of their birth cohort. We subsequently showed a survival advantage, mounting to 31%, for individuals with top 10% surviving first and second-degree relatives in both databases and across generations, even in the presence of non-longevous parents. To guide future genetic studies, we suggest to base case selection on top 10% survivors of their birth cohort with equally long-lived family members.

## Main

Human lifespan has a low heritability (12-25%)^1–3^, whereas survival into extreme ages (longevity) clusters within families^4–8^. Studies showed that parents, siblings^4–6,8–11^, and children^6, 12–16^ of longevous persons lived longer than first degree relatives of non-longevous persons or population controls. In addition, members of these longevous families seem to delay or even escape age-related diseases^17–20^ and in fact, healthy ageing in such families is marked by well attuned immune systems and metabolic health^21–23^. Understanding the genetic factors influencing longevity may provide novel insights into the mechanisms that promote health and minimize disease risk^1,24^. Identifying longevity loci, however, has been challenging and only a handful of genetic variants have been shown to associate with longevity across multiple independent studies^24–31^. The most consistent evidence has been obtained for variants in *APOE* and *FOXO3A* genes^24–29, 32^ in either genome-wide association studies (GWAS) or candidate gene studies.

The lack of consistent findings in longevity studies hampers comparative research and may be explained by genetic and environmental heterogeneity on one hand and uncertainty in defining the longevity trait itself, as illustrated by the large variation of longevity definitions on the other hand^1,3,6,9,12–16,18,19,24–31,33–37^. Establishing a threshold that best isolates the genetic component of longevity and including mortality information of family members is important because the environmentally-related increase in lifespan over recent decennia has caused an increase in longevity phenocopies. As a result, genetic longevity studies generally focus on singletons (i.e. individuals without longevous family members), selected based on one generation of mortality data^26,27,30,31,38^. Here, we aim to establish the threshold for longevity in unselected (for survival) multigenerational families and determine the importance of longevous family members for case selection so that those insights can be used in genetic studies to identify novel longevity loci.

We used the data available in the Utah Population Database (UPDB,Utah) and the LINKing System for historical family reconstruction (LINKS,Zeeland) based on US and Dutch citizens, respectively. Zeeland was a region with difficult living conditions compared to Utah (see methods section). In these datasets we identified 21,046 three-generational families (F1-F3) containing index persons (IPs, F2), their parents (F1), siblings (F2), spouses (F2), and children (F3) comprising 321,687 persons in total. First, we examined the association between the number of parents (F1) and siblings (F2) belonging to the top 1-60% of their birth cohort, in a cumulative way (comparing mutually inclusive percentile groups) with the survival of IPs (F2). Second, we determined the survival percentile threshold that drove the cumulative effects as a criterion for defining human longevity by investigating IPs (F2) who were divided into mutually exclusive groups based on the longevity of their parents (F4) and siblings (F2). Third, we focused on the top 10% parents and siblings to investigate whether longevous and non-longevous parents, with increasing numbers of longevous siblings, transmit longevity to the IPs. Fourth, we confirmed our findings in the next generation (F3) by examining the association between the longevity of IPs (F2), their spouses (parents, F2) and siblings (aunts and uncles, F2) with survival of IPs’ children (F3). Finally, we explored potential environmental influences by studying spouses (F2) of longevous IPs (F2).

## Results

We identified three generations of families in the UPDB and LINKS covering 10,929 and 10,117 families, respectively, who were centered around a single index person (IPs,F2) per family (Figure 1). We identified parents (F1, N_UPDB_=21,858 & N_LINKS_=202,343), siblings (F2, N_UPDB_=57,207 & N_LINKS_=53,999), spouses (F2, N_UPDB_=11,908 & N_LINKS_=10,791), and children (F3, N_UPDB_=62,145 & N_LINKS_=62,499) for all IPs in both datasets (Table 1). IPs were born between 1767 and 1929 in the UPDB, and between 1797 and 1908 in LINKS. In the UPDB, 52% of the IPs were female, compared to 53% in LINKS. The IPs mean age at death was 71.15 (SD=16.20) years in the UPDB and 63.85 (SD=17.99) years in LINKS. No IPs were censored, as they were selected to have an available birth and death date. In the following sections we explored associations between IP survival and the number of 1-60% surviving parents and siblings in a cumulative analysis and subsequently identified in mutually exclusive IP groups the survival percentile threshold that drives the cumulative effect and demarcates longevity (see methods section).

**Figure 1:**
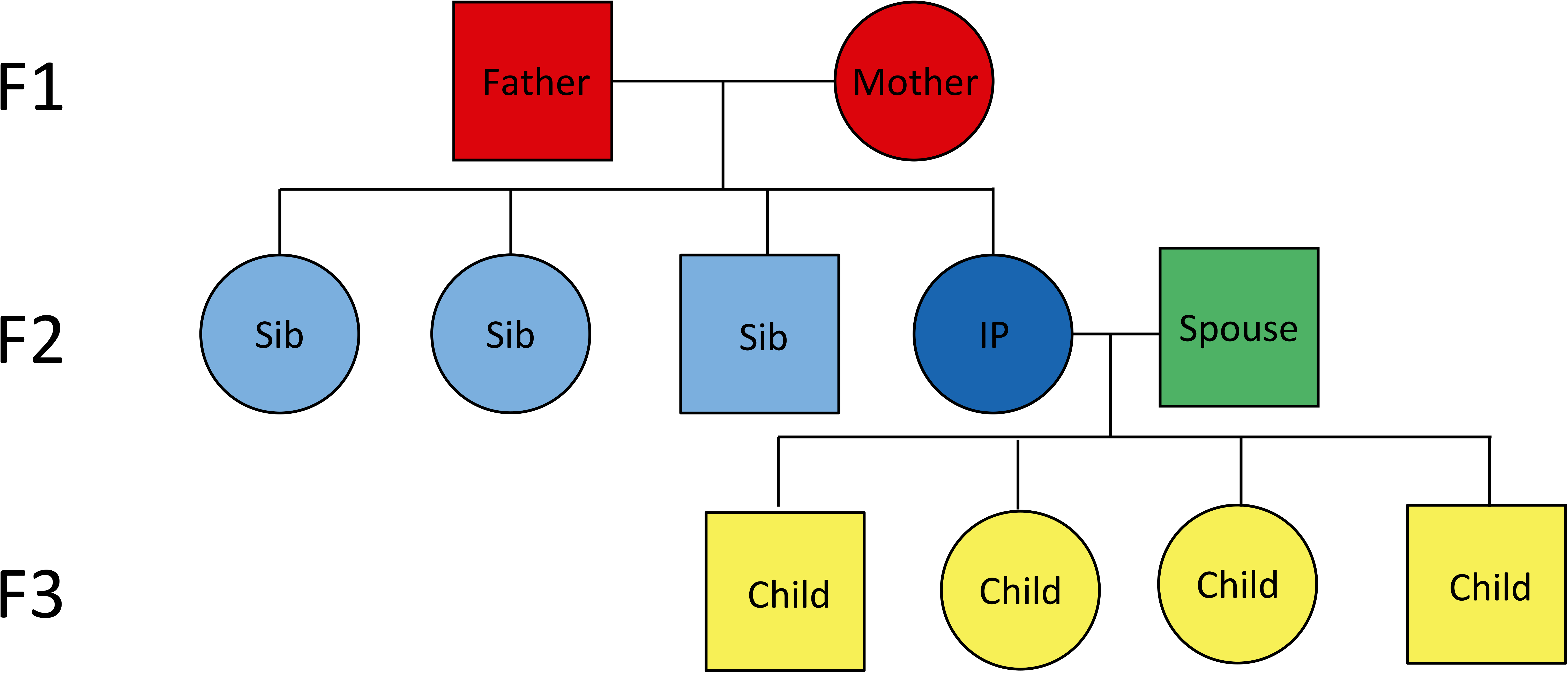
Conceptual pedigree of the 3 filial (F) generation families in the current study design. -This figure represents a hypothetical family from the UPDB or LINKS covering 3 filial (F) generations -Circles represent women, Squares represent men -DARK BLUE: Index persons ( F2), RED: Parents ( F1), LIGHT BLUE: Siblings of IP ( F2), GREEN: Spouses of IP (F2), YELLOW: Children of IP (F3). -IP: Index Person, Sib: Sibling, F: Filial.

**Table 1:** Overview of UPDB and LINKS Index Persons and their first degree relatives + spouses

| UPDB |  |  |  |  |  |
| --- | --- | --- | --- | --- | --- |
|  | Parents F0 | Index<br>persons F1 | Siblings F1 | Spouses F1 | Children F1 |
| **Number, N** | 21858 | 10929 | 57207 | 11908 | 62145 |
| **Deceased, N (%)** | 20453 (94) | 10929 (100) | 48300 (84) | 10863 (91) | 54797 (88) |
| **Female, N (%)** | 10929 (50) | 5641 (52) | 27627 (48) | 5978 (50) | 30151 (49) |
| **Range birth cohorts** | 1753 - 1906 | 1767 - 1929 | 1756 - 1932 | 1768 - 1929 | 1792 - 1954 |
| **Mean ad or al, years (SD)** | 68.80 (15.50) | 71.15 (16.20) | 44.85 (33.64) | 69.24 (17.63) | 54.95 (32.07) |
| **Mean ad, years (SD)** | 70.12 (15.20) | 71.15 (16.20) | 50.43 (32.50) | 70.92 (16.43) | 58.00 (31.27) |
| **Missing age, N (%)** | 437 (2) | 0 (0) | 835 (1) | 378 (3) | 365 (1) |
| **Censored, N (%)** | 968 (4) | 0 (0) | 8072 (14) | 667 (6) | 6983 (11) |
| LINKS |  |  |  |  |  |
|  | Parents F0 | Index<br>persons F1 | Siblings F1 | Spouses F1 | Children F1 |
| **Number, N** | 20234 | 10117 | 53999 | 10791 | 62499 |
| **Deceased, N (%)** | 15540 (77) | 10117 (100) | 40099 (74) | 8821 (81) | 43900 (70) |
| **Female, N (%)** | 10117 (50) | 5340 (53) | 25925 (48) | 5194 (48) | 30281 (49) |
| **Range birth cohorts** | 1740 – 1877 | 1797 - 1908 | 1796 – 1916 | 1775 - 1909 | 1818 – 1952 |
| **Mean ad or al, years (SD)** | 54.70 (20.00) | 63.85 (17.99) | 20.84 (27.98) | 59.03 (21.23) | 24.86 (30.86) |
| **Mean ad, years (SD)** | 62.65 (16.00) | 63.85 (17.99) | 23.94 (30.76) | 65.69 (17.21) | 29.59 (33.63) |
| **Missing age, N (%)** | 49 (<1) | 0 (0) | 14 (<1) | 27 (<1) | 21 (<1) |
| **Censored, N (%)** | 4645 (23) | 0 (0) | 13886 (26) | 1943 (18) | 18578 (30) |

-ad = age at death, al = age at last observation
-Missing age means that we have no observation at all.
-Number death refers to the number of deceased individuals .

### The number of top 1-60% parents and siblings strongly associates with the survival of IPs

For a first examination of the association between the number of parents (1 or 2,F1) and siblings (1 or 2+,F2) and IP (F2) survival and to explore if a larger level of family aggregation, in terms of numbers of parents (F2) and siblings (F2), was more evident at extreme survival percentiles, we fitted Cox regressions for each subsequent survival percentile (1^st^ to 60^th^ percentile). Figure 2A and C show that IPs with 1 parent belonging to the top 1-60%, had a survival advantage over IPs without a parent belonging to the top 1-60%. This was shown by the lowest observed statistically significant hazard ratios (HR) of 0.80 (95% CI_max-top 1%_=0.73-0.87) in the UPDB and 0.74 (95 CI_max-top 1%_=0.64-0.86) in LINKS where ‘max’ refers to the age with the largest effect. These HRs indicate a 20% and 26% lower hazard of dying respectively and from here we will refer to this as a 20% and 26% survival advantage. Having 2 parents belonging to the top 1-60% provided a stronger survival advantage to IPs (HR_max-top 2%-UPDB_=0.62 (95%CI=0.48-0.80) and HR_max-top 13%-LINKS_=0.75 (95% CI=0.61=0.93)), although Figure 2 shows that the power to detect survival effects of IPs with 2 longevous parents up to the 10^th^ percentile was weak for LINKS due to low group sizes.

**Figure 2:**
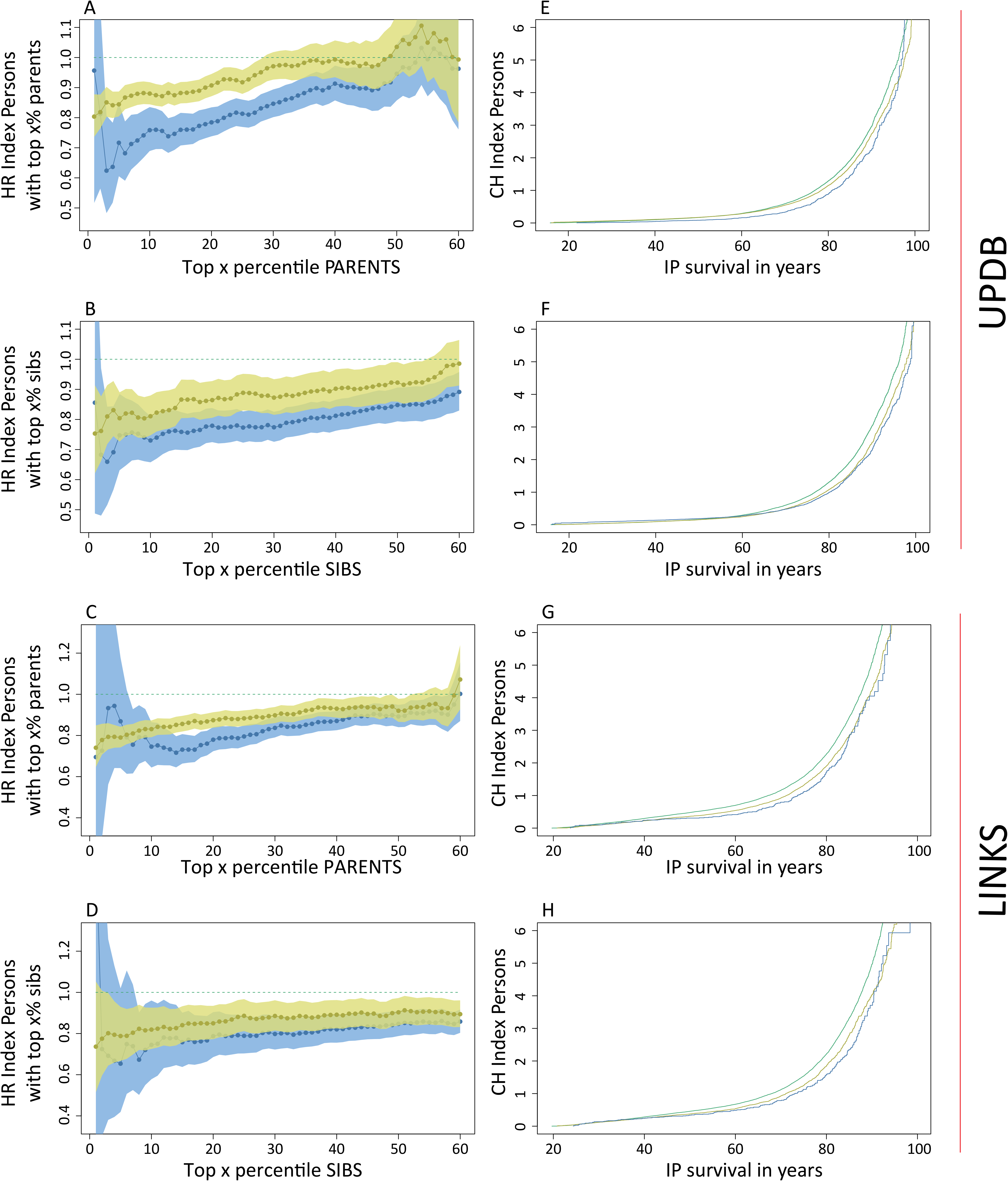
Survival of IPs with parents and siblings belonging to the 1^st^ until 60^th^ percentile survivors of their birth cohort. -This figure depicts the Hazard Ratio (HR) for IPs (left column) with 1 and 2 parents or 1 and 2+ siblings belonging to the top x percentile (x = 1,2,3, …, 60) of survivors of their birth cohort. The percentile groups (x-axis) are mutually inclusive, meaning that a first-degree family member who belonged to the top 1% also belonged to the top 5% etc. The figure also depicts the Cumulative Hazard (CH) for IPs (right column) with 1 and 2 parents or 1 and 2+ siblings who belong to the top 10%. -Green (dotted) lines present the reference group of 0 top x percentile parents or siblings, yellow lines represent 1 top x percentile parents or siblings, blue lines represent 2 or 2+ top x percentile siblings. -left column: x-axes represent the top x birth cohort based survival percentile, the y-axes represent the hazard ratio (HR) of dying for IPs having 1 and 2 or 2+ top x percentile parents or siblings compared to having 0 top x percentile parents or siblings. -right column: x-axes represent IP years of survival, y-axes represent the IPs’ cumulative hazard of dying while having 1 and 2 or 2+ top 10th percentile parents or siblings compared to having 0 top 10th percentile parents or siblings. -All estimates are adjusted for religion (UPDB only), sibship size, birth cohort, sex, socio-economic status, mother’s age at birth, birth order, birth intervals, twin birth, and number of top 10% parents or number of top 10% siblings for the sibling and parent analyses respectively.

The association of IP survival with longevous siblings was shown in Figure 2B and D. The maximum statistically significant HRs for IPs with 1 longevous sibling were 0.75 (95% CI_max-top 1%_=0.62-0.75) and 0.80 (95% CI_max-top 1%_=0.64-0.99) in the UPDB and LINKS respectively. For IPs with 2 or more longevous siblings these HRs were 0.66 (95% CI_max-top 3%-UPDB_=0.51-0.84) and 0.74 (95 CI_max-top 8%-LINKS_=0.55-0.99). The slopes in Figure 2A-D show a slight increase of IP survival advantage with the increase in percentile score. For example, IPs with parents with the best survival (the left most end of the x-axis) have lower hazard rates than IPs with the least survival (the right most end of the x-axis). We conclude that IP survival when expressed in HRs, both in UPDB and LINKS, increased with the number of longevous parents, with the number of longevous siblings and, though modestly, with the increase of parent and sibling survival percentile scores in a linear fashion as observed in Figure 2.

### Top 10-15% surviving family members demarcates the longevity effects of IPs

To determine the survival percentile threshold that drove the survival advantage of IPs (F2) with the number of top 1-60% parents (F1), we constructed 6 mutually exclusive IP (F2) groups (g) based on the survival of F1 parents (g1=[≥0^th^ & ≤1^th^ percentile], g2=[≥1^th^ & ≤5^th^ percentile], g3=[≥5^th^ & ≤10^th^ percentile], g4=[≥10^th^ & ≤15^th^ percentile], g5=[≥15^th^ & ≤20^th^ percentile], g6=[≥20^th^ & ≤100^th^ percentile],see methods section) and compared groups 1-5 with group 6. Figure 3A and B show the HRs of IP groups for the UPDB and LINKS. IPs in group 1, 2, 3, and 4 showed a significant survival advantage compared to group 6, with the lowest HR for group 1 in both the UPDB and LINKS (HR_max-UPDB_=0.76 (95% CI=0.67-0.86) and HR_max-LINKS_=0.71 (95% CI=0.59-0.86)). Group 5 did not statistically differ from group 6 (HR_group5-UPDB_=1 (95% CI=0.91-0.109) and HR_group5-LINKS_=0.96 (95% CI=0.87-1.05)) and thus, these effects indicate that the top 15% surviving parents drove the association with survival advantage of IPs as shown in Figure 2.

**Figure 3:**
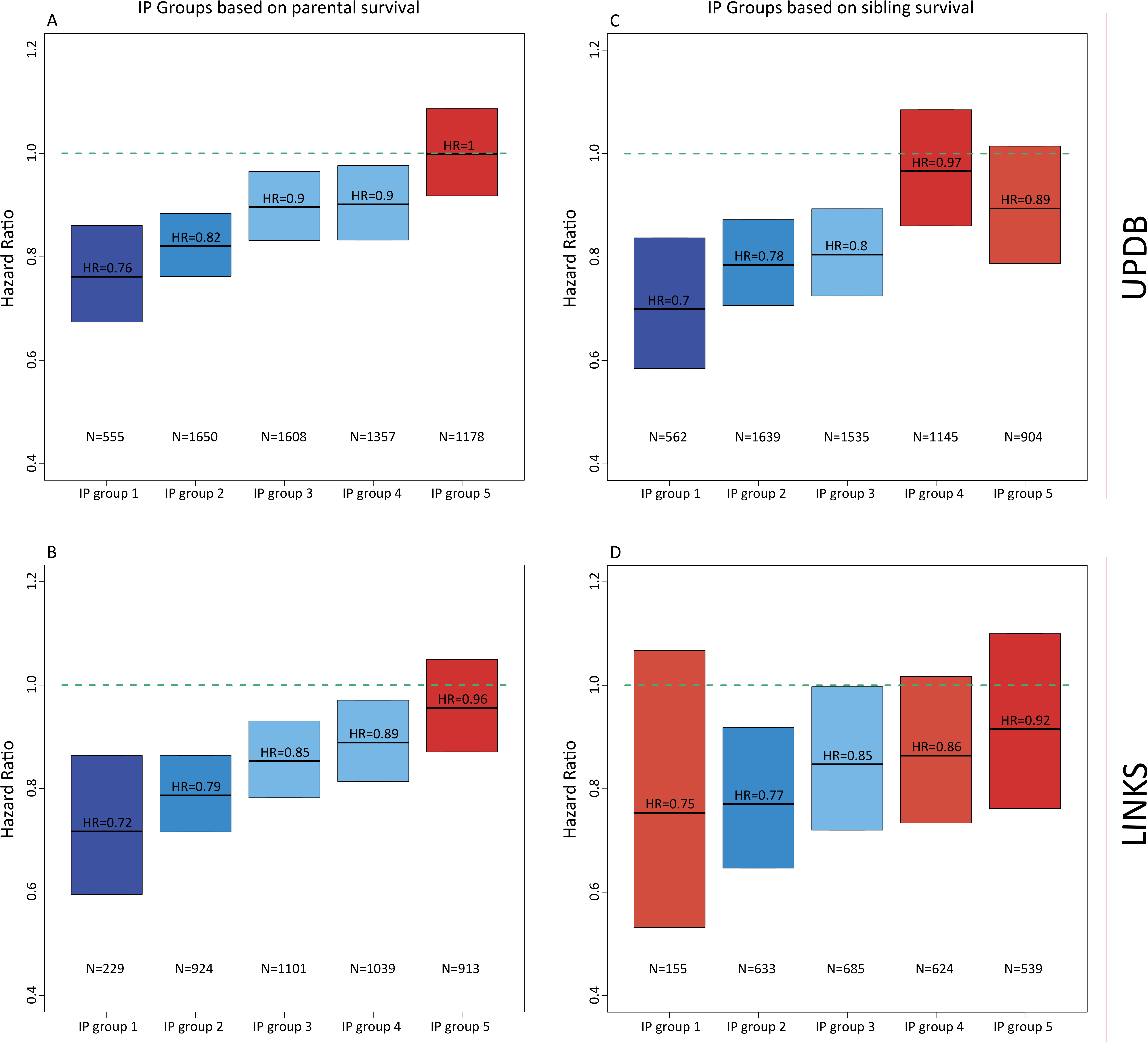
Hazard ratio for IPs grouped by their parental and sibling survival in mutual exclusive groups. -Parent and Sibling Groups: group 1 = IPs of whom the longest lived parent/sibling belonged to the [≥0th & ≤1th percentile] of their birth cohort, group 2 = IPs of whom the longest lived parent/sibling belonged to the [≥1th & ≤5th percentile], group 3 = IPs of whom the longest lived parent/sibling belonged to the [≥5th & ≤10th percentile], group 4 = IPs of whom the longest lived parent/sibling belonged to the [≥10th & ≤15th percentile], group 5 = IPs of whom the longest lived parent/sibling belonged to the [≥15th & ≤20th percentile], group 6 = IPs of whom the longest lived parent/sibling belonged to the [≥20th & ≤100th percentile]. -Groups were colored by the extremity of the HR. The darker the blue the stronger the survival benefit, the darker the red, the weaker the survival benefit and the effect was not significant in with the red colors. -The green lines represent the reference category, which is group 6. -N_green line_ at the top-right = 5144, N_green line_ at the top-left = 4581, N_green line_ at the bottom-right = 7481, N_green line_ at the bottom-left = 5911. -All estimates are adjusted for religion (UPDB only), sibship size, birth cohort, sex, socio-economic status, mother’s age at birth, birth order, birth intervals, twin birth, and number of top 10% parents or number of top 10% siblings for the sibling and parent analyses respectively.

In the same way we investigated the association of IPs (F2) survival with that of siblings (F2). Figure 3C and D show a survival advantage of IPs in UPDB group 1-3 and LINKS group 2 and 3 as compared to group 6 with the lowest HR for group 1 (UPDB) and group 2 (LINKS) (HR_group1-UPDB_=0.70 (95% CI=0.58-0.83) and HR_group2-LINKS_=0.77 (95% CI=0.64-0.92)), respectively. Group 4 and 5 did not significantly differ from group 6 (HR_group4-UPDB_=0.97 (95% CI=0.86-1.08) and HR_group4-LINKS_=0.86 (95% CI=0.73-1.02)) which indicated that both in the UPDB and LINKS the top 10% surviving siblings drove the association with the survival advantage of IPs as shown in Figure 2.

Based on the results presented in the cumulative and mutually exclusive group analyses we focused on the top 10% surviving family members because the mutually exclusive group analysis (analysis 2, Figure 3) indicated longevity effects up to the top 10% and 15% for siblings and parents respectively. Using the top 10% is consistent between the two groups and is a conservative choice. Furthermore, the cumulative analysis (analysis 1, Figure 2) indicated that the top 10% was a reasonable trade-off between effect size and group size (power) within and between the UPDB and LINKS. Hence, we explored the familial clustering of longevity and the influence of covariates for the top 10% surviving parents and siblings and verified all results in the subsequent generation. Next to the top 10% we also conducted our analyses on the top 5% which are illustrated in supplementary Figures 2-4 and supplementary Tables 5-9.

### The additive association between 10% surviving parents and siblings and the survival of IPs

Figure 2E-H show the cumulative hazard (CH) curves for IPs (F2) with 0, 1 and 2 or more, or exactly 2 parents/siblings (F1/F2) belonging to the top 10% of their birth cohorts and we show Kaplan-Meier and Nelson-Aalen baseline measures in supplementary Figure 5. Both in the UPDB and LINKS, the survival advantage associated with the number of top 10% siblings appears to start during the beginning (45 years in LINKS) and end (60 years in the UPDB) of the mid-life period. In both the UPDB and LINKS, the survival advantage of IPs with the number of top 10% parents started at the age of 35 years. It should be noted that early life effects could not be tested for because IPs were selected on having a child for the construction of three generation families.

Table 2 accompanies Figure 2E-H by showing the HRs for the number of top 10% parents (F1) and siblings (F2) and for the covariates we used to adjust the analyses. IPs with 1 top 10% parent had a maximum survival advantage of 13% and 17% compared to IPs without such a parent (HR=0.87_max-UPDB_ (95%CI=0.83-0.91) and HR=0.83_max-LINKS_ (95%CI=0.78-0.87)). The maximum statistically significant survival advantage for IPs with 2 top 10% parents was 27% and 29% (HR_max-UPDB_=0.73 (95%CI=0.65-0.79) and HR_max-LINKS_=0.71 95%CI=0.61-0.82)). The maximum statistically significant HR for having 1 top 10% sibling was 0.85 (95% CI_UPDB_=0.81-0.90) and 0.82 (95% CI_LINKS_=0.76-0.88). for 2+ top 10% siblings the HR was 0.75 (95% CI_UPDB_=0.67-0.83) and 0.81 (95% CI_LINKS_=0.65-1.02). The survival advantage of IPs with 1 and 2 or more, or exactly 2 top 10% siblings and parents respectively was independent of covariates such as sibship size and religion (LDS church). Religious IPs from Utah had a lower HR than non-religious persons (HR_UPDB_=0.72 (95% CI=0.65-0.79)) and in the UPDB we observed that sibship size had a small influence on the survival of IPs (HR_UPDB_=1.01 (95% Ci=1.00-1.02) whereas in LINKS sibship size had no significant effect HR_LINKS_=1.01 (95% CI=1.00-1.01)). The survival of IPs increased with the increase of birth cohort (HR_UPDB and LINKS_=0.99 (95% CI=[>0.99<1.00])) and women had a better survival than men, in the UPDB (HR_UPDB_=0.68 (95% Ci=0.64-0.72)) but not in LINKS (HR_LINKS_=1.03 (95% Ci=0.98-1.07)). Furthermore, In Utah, high socio-economic status IPs outlived low socio-economic status IPs whereas this was not the case in LINKS. The association between the number of longevous siblings/parents and the survival of IPs were independent of each other and no other statistically significant effect was observed for having both longevous parents and siblings. Moreover, the number of longevous siblings showed a strong association with the survival of IPs when both parents were non-longevous. The HR for 1 longevous sibling was 0.85 (95% CI=0.79-0.91) and the HR for 2 or more longevous siblings was 0.75 (95% CI=0.65-0.87) in the UPDB. The HR for 1 longevous sibling was 0.78 (95% CI=0.72-0.85) and the HR for 2 or more longevous siblings was 0.72 (95% CI=0.53-0.99) in LINKS (supplementary Table 2). In a final step, we observed no evidence that the association of IPs and parental survival depended on maternal or paternal effects, for example through transmission preferentially via the mother or father. (supplementary Table 3).

**Table 2:** Survival analysis for IPs by top 10% siblings and top 10% parents

|  | UPDB |  |  | LINKS |  |  |
| --- | --- | --- | --- | --- | --- | --- |
|  | N (mean) | HR (95% CI) | p-value | N (mean) | HR (95% CI) | p-value |
| **Top 10% parents (F1)** |  |  |  |  |  |  |
| 0 (ref) | 7129 (0.66) |  |  | 7864 (0.78) |  |  |
| 1 | 3345 (0.30) | 0.87 (0.83-0.91) | 3.79*10<sup>-9</sup> | 2096 (0.20) | 0.83 (0.78-0.87) | 2.40*10<sup>-13</sup> |
| 2 | 455 (0.4) | 0.73 (0.65-0.79) | 1.55*10<sup>-8</sup> | 184 (0.2) | 0.71 (0.61-0.84) | 3.42*10<sup>-5</sup> |
| **Top 10% sibs (F2)** |  |  |  |  |  |  |
| 0 (ref) | 7203 (0.66) |  |  | 8647 (0.85) |  |  |
| 1 | 2644 (0.24) | 0.85 (0.81-0.90) | 4.34*10<sup>-9</sup> | 1256 (0.13) | 0.82 (0.76-0.88) | 1.47*10<sup>-7</sup> |
| 2+ | 1082 (0.10) | 0.75 (0.67-0.83) | 1.77*10<sup>-8</sup> | 214 (0.2) | 0.81 (0.65-1.02) | 7.30*10<sup>-2</sup> |
| **LDS (F2)** |  |  |  |  |  |  |
| 0 - non-religious(ref) | 3101 (0.27) |  |  |  |  |  |
| 1 - baptized | 590 (0.06) | 0.72 (0.65-0.79) | 2.86*10<sup>-11</sup> | NA | NA | NA |
| 2 - baptized + |  |  |  |  |  |  |
| endowment | 6990 (0.65) | 0.84 (0.80-0.88) | 3.97*10<sup>-12</sup> | NA | NA | NA |
| 3 - missing | 248 (0.02) | 0.90 (0.78-1.04) | 1.43*10<sup>-1</sup> | NA | NA | NA |
| **Sibship size (F2)** | 10929 (6.22) | 1.01 (1.00-1.02) | 6.00*10<sup>-3</sup> | 10117 (6.35) | 1.01 (1.00-1.01) | 2.05*10<sup>-1</sup> |
| **Birth cohort, years (F2)** | 10929 (1870) | 0.99 (>0.99<1.00) | <1.00*10<sup>-15</sup> | 10117 (1835) | 0.99 (>0.99<1.00) | <1.00*10<sup>-15</sup> |
| **Sex (F2)** |  |  |  |  |  |  |
| Man (ref) | 5288 (0.46) |  |  | 4777 (0.48) |  |  |
| Women | 5641 (0.53) | 0.68 (0.64-0.72) | <1.00*10<sup>-15</sup> | 5340 (0.52) | 1.03 (0.98-1.07) | 2.60*10<sup>-1</sup> |
| **SES – OCC_1950 (F2)** |  |  |  |  |  |  |
| 0 - High (ref) | 345 (0.03) |  |  | 67 (0.01) |  |  |
| 1 | 1493 (0.14) | 1.11 (0.98-1.27) | 1.09*10<sup>-1</sup> | 645 (0.06) | 0.88 (0.68-1.14) | 3.30*10<sup>-1</sup> |
| 2 | 447 (0.04) | 1.16 (1.00-1.35) | 5.65*10<sup>-2</sup> | 536 (0.05) | 0.96 (0.74-1.24) | 7.63*10<sup>-1</sup> |
| 3 | 403 (0.04) | 1.29 (1.10-1.50) | 1.54*10<sup>-3</sup> | 62 (0.01) | 0.78 (0.55-1.11) | 1.74*10<sup>-1</sup> |
| 4 | 200 (0.02) | 1.15 (0.96-1.38) | 1.52*10<sup>-1</sup> | 71 (0.01) | 0.96 (0.68-1.36) | 8.18*10<sup>-1</sup> |
| 5 | 953 (0.09) | 1.28 (1.11-1.46) | 4.15*10<sup>-4</sup> | 733 (0.07) | 0.79 (0.61-1.02) | 7.14*10<sup>-2</sup> |
| 6 | 727 (0.07) | 1.34 (1.17-1.54) | 2.91*10<sup>-5</sup> | 311 (0.03) | 0.85 (0.65-1.11) | 2.42*10<sup>-1</sup> |
| 7 | 564 (0.06) | 1.30 (1.12-1.51) | 3.37*10<sup>-4</sup> | 759 (0.08) | 0.81 (0.62-1.04) | 1.03*10<sup>-1</sup> |
| 8 | 170 (0.01) | 1.24 (1.01-1.52) | 3.61*10<sup>-2</sup> | 575 (0.06) | 0.84 (0.65-1.08) | 1.78*10<sup>-1</sup> |
| 9 – Low | 588 (0.05) | 1.39 (1.21-1.61) | 7.38*10<sup>-6</sup> | 3656 (0.36) | 0.82 (0.64-1.05) | 1.10*10<sup>-1</sup> |
| 999 - missing | 5039 (0.45) | 1.58 (1.40-1.79) | 1.88*10<sup>-13</sup> | 2702 (0.26) | 0.91 (0.71-1.17) | 4.51*10<sup>-1</sup> |
| **Log likelihood** | -74984 |  |  | -77371 |  |  |

-Table corresponds to the CH curves in the top and bottom right panel of figure 2
-Means represent a mean for a continuous variable and a proportion for a categorical variable
-Additional covariates are: age mom at birth, birth order, birth intervals (in years), twin birth
-When the p-value was lower than 1.00e-15 we indicated the P-value as <1.00\*10<sup>-15</sup>
-LDS: the church of Jesus Christ of latter-day saints (Mormon church), SES: socio-economic status, OCC: occupational coding scheme of 1950

### Survival advantage for children with longevous parents and longevous aunts and uncles

We explored the robustness of our findings in F1 and F2 by examining the association between the longevity of IPs (F2), their spouses (F2) and siblings (F2) and the survival of IPs’ children (F2). We investigated whether longevity was transmitted from IPs (F2) to their children (F3) and if the children (F3) with longevous aunts and uncles (siblings of the IPs,F2) had a survival advantage compared to children (F3) without longevous aunts and uncles (F2). To test this, we fitted Cox regressions, with a random effect (frailty) to adjust for within-family relations of the F3 children. Table 3 shows that children of a top 10% surviving IP had a HR of 0.86 (95% CI_UPDB_=0.84-0.89) in the UPDB and 0.85 in LINKS (95% CI_LINKS_=0.82-0.88) compared to children without a top 10% IP. Moreover, results indicated that children with two top 10% parents (IPs and spouses) had a HR of 0.76 (95% CI_UPDB_=0.67-0.85) in the UPDB and 0.77 (95% CI_LINKS_=0.71-0.84) in LINKS. Similar to the IPs, we observed that the survival of children did not depend on maternal or paternal effects (supplementary Table 3).

**Table 3:** Frailty survival analysis for Children of IPs by top 10% IP’s and aunt and uncles of children

|  | UPDB |  |  | LINKS |  |  |
| --- | --- | --- | --- | --- | --- | --- |
|  | N (mean) | HR (95% CI) | p-value | N (mean) | HR (95% CI) | p-value |
| **Top 10% IP (F2)** |  |  |  |  |  |  |
| 0 non LL (ref.) | 49460 (0.80) |  |  | 53382 (0.85) |  |  |
| 1 LL | 12320 (0.20) | 0.86 (0.84-0.89) | <1.00*10<sup>-15</sup> | 9096 (0.15) | 0.85 (0.82-0.88) | <1.00*10<sup>-15</sup> |
| **Top 10% aunts and uncles (F2)** |  |  |  |  |  |  |
| 0 (ref.) | 40156 (0.65) |  |  | 53232 (0.85) |  |  |
| 1 | 15361 (0.25) | 0.96 (0.93-0.99) | 2.28*10<sup>-3</sup> | 7817 (0.12) | 0.96 (0.92-0.99) | 1.97*10<sup>-2</sup> |
| 2+ | 6263 (0.10) | 0.92 (0.88-0.96) | 2.45*10<sup>-5</sup> | 1429 (0.3) | 0.84 (0.78-0.92) | 5.56*10<sup>-5</sup> |
| **Sibshpsize (F3)** | 61780 (8.05) | 1.02 (1.01-1.02) | 8.77*10<sup>-15</sup> | 62478 (0.8.04) | 0.99 (0.99-1.00) | 8.50*10<sup>-1</sup> |
| **Birth year (F3)** | 61780 (1893) | 0.99 (>0.99<1.00) | <1.00*10<sup>-15</sup> | 62478 (0.1865) | 0.99 (>0.99<1.00) | 5.19*10<sup>-11</sup> |
| **Sex (F3)** |  |  |  |  |  |  |
| Man (ref.) | 31794 (51) |  |  | 32137 (0.52) |  |  |
| Women | 29976 (49) | 0.62 (0.61-0.63) | <1.00*10<sup>-15</sup> | 30272 (0.48) | 0.64 (0.63-0.66) | <1.00*10<sup>-15</sup> |
| **Famid intercept (variance)** | 0.34 (0.12) |  |  | 0.34 (0.11) |  |  |
| **BIC** | -24407.48 |  |  | -21514.91 |  |  |

-Additional covariates are: birth order, birth intervals (years), age mom at birth -Religion, Socio-economic status, twin birth have been stratified -When the p-value was lower than 1.00e-15 we indicated the P-value as <1.00e-15 -BIC: Bayesian Information Criterion -Famid: family identifier

Children with 1 or more top 10% aunts or uncles had a 4-19% survival advantage compared to children without such aunts or uncles (HR_min-UPDB_=0.96 (95% CI=0.93-0.99) and HR_max-LINKS_=0.84 (95% CI=0.78-0.92)) and this effect was independent of having a top 10% parent (either the IP or the IP’s spouse). A stratified analysis showed that the survival benefit for children with the number of top 10% aunts and uncles was still strongly present when the IP and the IP’s spouse were non-longevous (HR_min-UPDB - 1 aunt/uncle_=0.96 (95% CI=0.93-0.99) and HR_max-LINKS - 2+ aunts/uncles_=0.81 (95% CI=0.73-0.90)) (supplementary Table 4). Lastly, supplementary Figure 6 shows that the survival benefit for children of a longevous IP and a longevous IP with a longevous spouse (i.e. 1 or 2 longevous parents) started from birth (LINKS) and very early in life (UPDB).

### Spouses live longer in Zeeland but not in Utah

Familial clustering of longevity may depend on (later life) shared environmental effects which could also provide survival benefits to the spouses (F2) of longevous IPs (F1). Hence, we divided the spouses (F2) into mutually exclusive groups according to the survival of the IPs (see methods). Figure 4A and 4C show that none of the spouse groups in the UPDB differed from reference group 6 or from any of the other groups, indicating no survival benefit for spouses. In LINKS (Figure 4B and 4D), spouses of IPs with the highest survival percentile (group 2) had a 15% (HR_group2-LINKS_=0.85 (95% CI=0.78-0.93)) survival advantage compared to group 6 spouses. This survival advantage was similar for spouses of IPs in group 3,4, and 5 (HR_group3-LINKS_=0.86 (95% CI=0.79-0.93); HR_group4-LINKS_=0.92 (95% CI=0.85-0.99); HR_group5-LINKS_=0.86 (95%CI=0.79-0.93)). For Group 1 the effect was comparable but not significant (HR_group1-LINKS_=0.85 (95% CI=0.70-1.04)), the test in group 4 did not meet Bonferroni correction for multiple testing.

**Figure 4:**
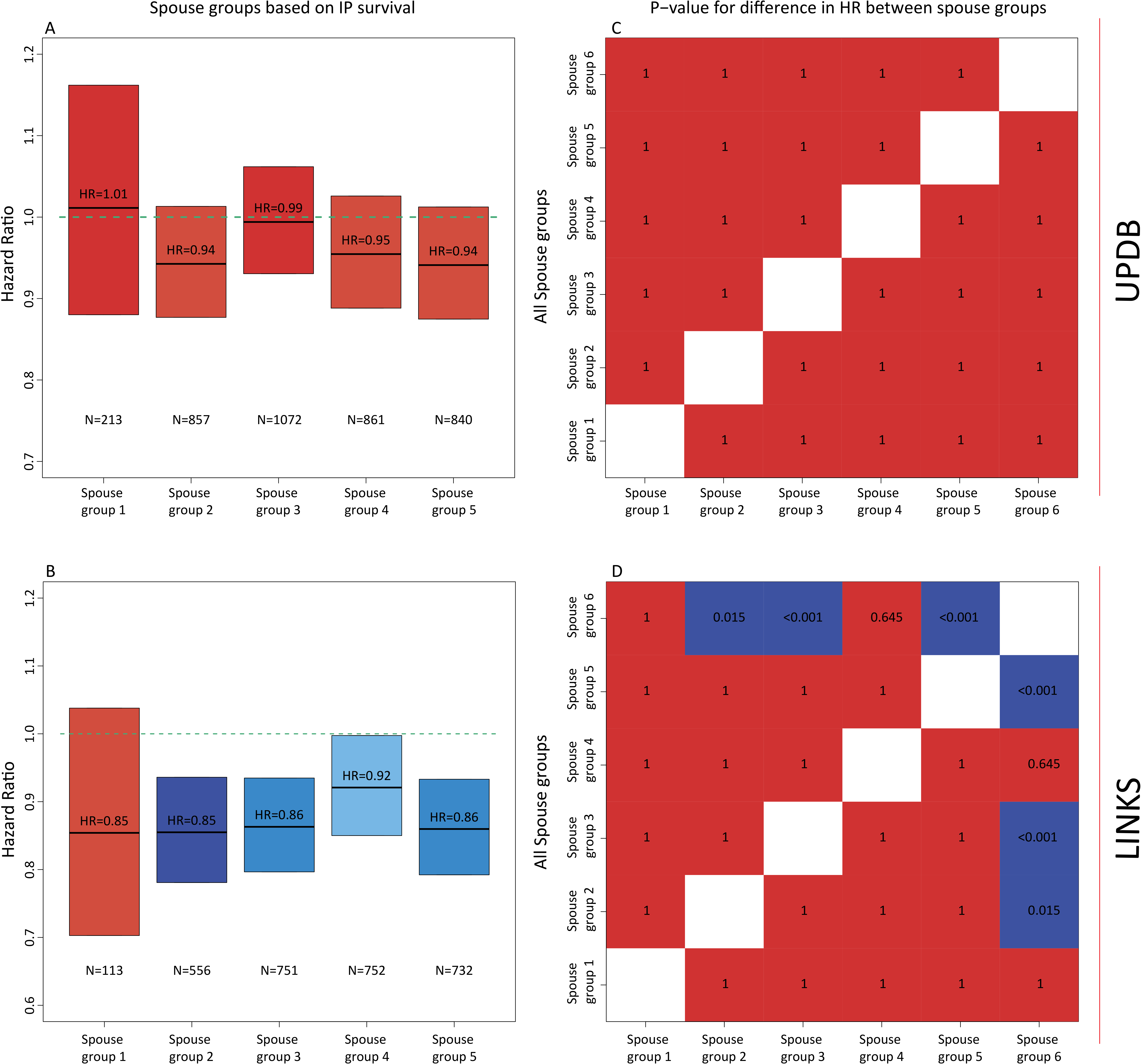
Hazard ratio for spouses grouped by the survival of their IP in mutual exclusive groups. -Spouse Groups: group 1 = Spouses of whom the IP belonged to the [≥0th & ≤1th percentile] of their birth cohort, group 2 = Spouses of whom the IP belonged to the [≥1th & ≤5th percentile] of their birth cohort, group 3 = Spouses of whom the IP belonged to the [≥5th & ≤10th percentile] of their birth cohort, group 4 = Spouses of whom the IP belonged to the [≥10th & ≤15th percentile] of their birth cohort, group 5 = Spouses of whom the IP belonged to the [≥15th & ≤20th percentile] of their birth cohort, group 6 = Spouses of whom the IP belonged to the [≥20th & ≤100th percentile] of their birth cohort. -Groups were colored by the extremity of the HR. The darker the blue the stronger the survival benefit, the darker the red, the weaker the survival benefit and the effect was not significant in with the red colors. -The green lines represent the reference category, which is group 6. -N_green line_ at the top-left = 8065, N_green line_ at the bottom-left = 7887, -The right column represents a post-hoc test of all groups and illustrates the p-values for the differences in HR between the spouse groups. Blue color indicates a statistically significant effect after bonferroni correction, red color indicates a non-statistically significant effect after bonferroni correction. -All estimates are adjusted for religion (UPDB only), sibship size, birth cohort, sex, socio-economic status, mother’s age at birth, birth order, birth intervals, twin birth, and number of top 10% parents or number of top 10% siblings for the sibling and parent analyses respectively.

## Discussion

Human longevity clusters within specific families. Insight into this clustering is important, especially to improve our understanding of genetic and environmental factors driving healthy aging and longevity. The analyses of the UPDB and LINKS datasets provide strong evidence that for longevous (up to the top 10%) survivors and their families, longevity is transmitted as a quantitative genetic trait. The main observations supporting this notion are (1) in both datasets the survival of F2 IPs (and their F3 children) increased with each additional longevous parent (F1 and F2) and sibling (2), (2) in both datasets the survival of IPs (F2) increased with the number of longevous siblings (F2) in the absence of longevous parents (F1) and likewise the survival of IPs’ children (F3) increased with the number of longevous aunts and uncles in the absence of longevous parents.

Longevity was transmitted even if parents themselves did not become longevous, which supports the notion that a beneficial genetic component was transmitted. In addition, children of non-longevous parents. Further evidence for the transmission of a genetic component was shown by the fact that none of the tested environmental confounders affected the associations between parental/sibling longevity and IP/children survival. In addition, the fact that we observed very similar results between the two databases, which cover populations with inherently different environmentally related mortality regimes, significantly adds to the robustness of our observations regarding the associations between parental/sibling longevity and IP (F2) and children (F3) survival.

We showed that spouses (F2) who married longevous IPs (F2) did not live significantly longer than spouses (F2) who married a non-longevous IP (F2) in the UPDB while they did in LINKS. Literature is inconclusive about the potential survival advantage of spouses of long-lived persons^5,6,8,39,40^. Pedersen et al. (2017) observed a survival advantage in the Long Life Family Study for spouses of long-lived siblings when comparing them to a birth cohort and sex matched control group. The authors point to assortative mating as a factor explaining the survival advantage for these spouses^5^. A Quebec study, focused on the spouses of 806 centenarians, also reported a survival advantage^39^ and a study of Southern Italy demonstrated that male nonagenarians outlived their spouses, whereas this was not the case for female nonagenarians^40^. A recent study showed that the spouses of 944 nonagenarians had no survival advantage but a life-long sustained survival pattern similar to the general population^8^. An explanation for the difference between the UPDB and LINKS datasets may possibly be that Zeeland had a higher level of relatedness than in Utah. Zeeland had poor living conditions^41^ and was characterized by out migration to other provinces or abroad, but limited mobility within the province to other places^42^. Utah at that time had better living conditions^43^ with continuous streams of freshly incoming migrants, ensuring a steady influx of new genes^44^, creating high genetic diversity. Hence, it could be that in Zeeland, spouses and IPs were often related to each other and thus shared some of the genetic component contributing to longevity.

In all our analyses, except for the spouse analysis, we adjusted for religion (UPDB only), sibship size, birth cohort, sex, socio-economic status, mother’s age at birth, birth order, birth intervals, and twin birth. Some of these biological, social, and demographic factors associated with the mortality of IPs (F2) and their children (F3). Nevertheless, these covariates neither confounded the association between parental (F1) and sibling (F2) longevity and IP (F2) survival, nor that between IP (F2) and spouse (F2) longevity and their children’s (F3) survival or between longevity of aunts and uncles (F2) and the survival of IPs’ children (F3). We, however, cannot rule out that other, unobserved non-genetic familial effects affect our results. Furthermore, using either Swedish or Dutch lifetables to determine survival percentiles was quite strict for Zeeland because of the hazardous environment^41^. As a result, the number of longevous persons was quite low in LINKS relative to the UPDB. Although the IPs were randomly selected, we could not completely rule out selection effects, for example related to early life mortality. However, confirmation of the F1-F2 results in the next generation F2/F3 significantly strengthens the results and allowed us to cope with the potential selection effects for IPs. Unlike observations we previously made in the Leiden Longevity Study^8^ concerning maternal effects on longevity in the generation of the nonagenarians and their parents, we did not observe evidence for a stronger transmission from either parent to the IPs (F1 to F2), or from IPs to their children (F2 to F3) in our current study. We cannot draw final conclusions on this aspect because for the F1-2 transmission we may have missed parental influences on early life mortality since IPs were selected for having survived to an age at which they had one child. However, we did capture early life mortality for F2-F3 but in those generations the selection pressure on child mortality was already slightly decreasing^45^.

Although lifespan is not very heritable in the population at large^3^ recent studies have been able to identify^30,31^ and replicate^46^ some lifespan associated alleles that lower the risk of age related diseases. Our results imply that to find loci that promote survival to the highest ages in the population, genetic studies should be based on long lived cases including at least parental mortality information but preferably also mortality information of siblings and other first and second degree family members. The longevity threshold should include cases belonging up to the top 10% survivors, with parents belonging up to the top 15% survivors of their birth cohort and siblings belonging up to the top 10% survivors of their birth cohort. To increase the longevity effect, the percentile threshold applied may even be more extreme but would likely lead unnecessarily to an underpowered design. If we consistently apply our suggested longevity definition across studies we may improve the comparative nature of longevity studies and create a new impulse to detect novel genetic variants.

## Methods

### Population datasets

#### Utah Population Database

The Utah Population Database (UPDB) contains demographic and genealogical information which is linked to medical records. The data construction began in the mid-1970s with genealogy records from the archives at the Utah Family History Library and was initially based on the founding members of the Utah population, their descendants, and then subsequently all individuals living in Utah. These records contain demographic and mortality information on the pioneers of Utah (United States), their parents and children, and have been linked into multigenerational pedigrees. The founding families were selected for the UPDB when at least one member had a vital event (birth, marriage, or death) on the Mormon pioneer trail or in Utah. The UPDB has been expanded to incorporate other high-quality, state-wide data sources, such as birth and death certificates, cancer records, driver license records, and census records. Currently the UPDB contains information on more than 11 million individuals and covers a maximum of 17 generations^47,48^.

#### LINKing System for historical family reconstruction

The LINKing System for historical family reconstruction (LINKS) data contains demographic and genealogical information which was derived from linked vital event registers (birth, marriage, and death certificates). The data indexing began in 1995 by the “*Zeeuws*” archive and the results were published by way of “WieWasWie”. The data currently covers over 25 million Dutch vital event records^49,50^. Data construction has been completed for the province of Zeeland and is still ongoing for the other provinces in the Netherlands. Currently LINKS Zeeland (henceforth referred to as LINKS) contains 739,453 birth, 387102 marriage, and 641,216 death certificates which were linked together to reconstruct intergenerational pedigrees and individual life courses^42^. In total the Zeeland data contains 1,930,157 persons covering a maximum of 7 generations^51^.

#### Historical context in of Utah and Zeeland

Both Utah and Zeeland were high fertility populations^41,43,52^, with a mean number of children of around 7 during the period of this study (1740-1954). In general, Utah was marked by healthy living conditions and Zeeland by contrast, was a much unhealthier place to live. One of the main reasons for the unhealthy living conditions in Zeeland was the lack of clean drinking water, the high prevalence of waterborne diseases and of malaria^41,53,54^. In Utah the quality of the drinking water was good, since water from melting snow, that was filtered running of the mountains, was used to drink^43^. The differences in living conditions between Utah and Zeeland were reflected by a relatively low infant and childhood mortality in Utah^55^ and high mortality rates for infants and children in Zeeland^54^, especially before 1900. Moreover, Utah was known to be a high in-migration population^44^ whereas there were indications that Zeeland had a low influx and outflux of migrants^42^.

#### Study selection

For the current study we used 3 Filial (F) generations (F1-3) from the UPDB and LINKS (N_UPDB+LINKS_=321,687). We reconstructed families in both datasets and denote generation 1 as the starting point of the pedigrees in the data. The starting point for this study was generation 3 because starting here minimized missing family links and birth or death dates due to the nature of the source material underlying the data. We denote generation 3 as filial generation 1 (F1). Subsequently, the children (N_UPDB+LINKS_=132,247) of the F1 parents were identified (F2) so that unique families were represented by 2 parents (F1) and their offspring (F2). Next an index person (IP,F2) was randomly selected per F2 sibship (N=21,046) meeting the following criteria: (1) The date of birth and death had to be available, (2) At least one child, sibling, and spouse had to be available, (3) sex had to be available, (4) for the UPDB data only, the IP should preferably be identifiable on a genealogy record (supplementary Table 1). From there we identified the siblings (F2, N_UPDB+LINKS_=88,399), spouses (F2, N_UPDB+LINKS_=22,699), and the children (F3,N_UPDB+LINKS_=124,644) of the IPs (Table 1 and Figure 1). To summarize, both in the UPDB and LINKS we identified IPs (F2), their parents (F1), siblings (F2), spouses (F2), and children (F3).

### Lifetables

We used cohort lifetables to calculate birth cohort and sex specific survival percentiles for each individual in the UPDB and LINKS. This approach prevents against the effects of secular mortality trends over the last centuries and enables comparisons across study populations^1,11^. We could not use United States (US) lifetables because cohort lifetables were not available at all and period lifetables were only available from 1933 onward. However, for Sweden and the Netherlands, population based cohort lifetables were available from 1751 and 1850 until 2018 respectively^56–59^. These lifetables contained, for each birth year and sex, an estimate of the hazard of dying between ages x and x + n (h_x_) based on yearly intervals (n=1) up to 99 years of age. Conditional cumulative hazards (H_x_) and survival probabilities (S_x_) were derived using these hazards. In turn, we could determine the sex and birth year specific survival percentile for each person in our study. Swedish cohort lifetables date back furthest of all available lifetables and were shown to be consistent with the lifetables of multiple industrialized societies^60^. In addition, we ensured that the survival percentiles were calculated in the same way for the UPDB and LINKS to make a fair comparison between the survival percentiles. Hence, the Swedish cohort lifetables were used for both datasets and for the LINKS data the Dutch lifetables were used as a sensitivity analysis. Supplementary Figure 1 shows the ages at death corresponding to the top 10, 5, and 1 percent survivors for the UPDB and LINKS and can be used to map the percentiles to absolute ages.

### Statistical analyses

Statistical analyses were conducted using R version 3.3.0^61^. We reported 95% confidence intervals (CIs) and considered p-values statistically significant at the 5% level (α = 0.05).

#### Exploring the association between the number of parents and siblings with IP survival at increasingly extreme survival percentiles (analysis 1)

To determine if (1) the association between the survival of IPs and the survival of their parents and siblings increased with increasing survival percentiles, and (2) a larger level of family aggregation, in terms of numbers of parents and siblings, was more evident at extreme survival percentiles, we investigated the association between the IP survival and the number of parents and siblings reaching increasingly more extreme survival percentiles. We sequentially identified the number of parents and siblings belonging to the top x (x = 1,2,3, …, 60) percentiles of their birth cohorts (from here: percentiles) and we analyzed their association with the survival of the IPs for each subsequent percentile using a Cox proportional hazard model:

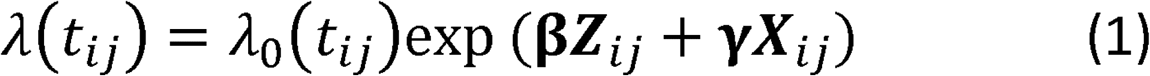

where *t_ij_* is the age at death or the age at last follow-up for IP *j* in family *i*. λ_0_(*t_ij_*) refers to the baseline hazard, which is left unspecified in a Cox-type model. **β** is the vector of regression coefficients for the main effects of interest (***Z***) which correspond to: (1) the number of parents belonging to the top x percentile, (2) and the number of siblings belonging to the top x percentile. **γ** is a vector of regression coefficients for the effects of covariates and possible confounders (***X***) which are: IPs’ religion (UPDB only), sibship size, birth cohort, sex, socio-economic status, mother’s age at birth, birth order, birth intervals, and twin birth.

#### Identifying a survival percentile threshold that demarcates longevity (analysis 2)

The previous analysis, bases on the cumulative effects, does not allow us to identify a specific threshold to define longevity, since the top x percentiles were not mutually exclusive, i.e., if a person belonged to the top 1% survivors, this person also belonged to the groups of top 5% and top 10% survivors. To determine the survival percentile threshold that drove the cumulative top x percentile effects described in the previous section, we grouped IPs according to the survival of their parents and siblings for two separate analysis. More specifically, we constructed mutually exclusive groups of IPs based on having at least one parent or sibling belonging to group g (g = 1,2,3, …, 6): group 1 = [≥0^th^ & ≤1^th^ percentile], group 2 = [≥1^th^ & ≤5^th^ percentile], group 3 = [≥5^th^ & ≤10^th^ percentile], group 4 = [≥10^th^ & ≤15^th^ percentile], group 5 =[≥15^th^ & ≤20^th^ percentile], group 6 = [≥20^th^ & ≤100^th^ percentile]. Group membership was defined by the most long-lived parent or sibling of the IP. Using Cox proportional hazards models (see expression (1)), we compared the effects of all groups to reference group 6, corresponding to IPs with all parents or siblings belonging to the 20^th^ or less extreme survival percentile and multiple combinations of defining group 6 were tested. Here, the **β** is the vector of regression coefficients for the main effects of interest (***Z***) which correspond to: (1) the IPs who were divided into mutually exclusive groups by their parental mortality and (2) the IPs who were independently grouped by their sibling mortality. Other parts of the expression are the same as noted in expression 1.

#### Exploring the top 10% parents, siblings, and covariates in an integrated design (analysis 3)

Based on the analyses expressed in the previous section we chose the top 10% survivors for specific follow-up analyses. Based on the results presented in the cumulative and mutually exclusive group analyses we focused on the top 10 surviving family members because the mutually exclusive group analysis (analysis 2) indicated longevity effects for siblings beyond the top 10% and 15% for siblings and parents respectively. Using the top 10% is consistent between the two groups and is a conservative choice. Furthermore, the cumulative analysis (analysis 1) indicated that the top 10% was a good trade-off between effect size and group size (power) within and between the UPDB and LINKS. Hence, we focused on top 10% parents and siblings in an integrated design to investigate the association between IP survival and the number of parents and siblings belonging to the top 10%. We subsequently investigated the association between the number of top 10% siblings and IP survival for IPs without top 10% parents, using Cox regression (see expression (1)). Here the **β** is the vector of regression coefficients for the main effects of interest (***Z***) which correspond to: (1) the number of parents and siblings belonging to the top 10% and (2) the number of siblings belonging to the top 10% for IPs without top 10% parents. Other parts of the expression are the same as noted in expression 1.

In all Cox regression analyses, based on expression 1, we accounted for the fact that IPs were selected to have a spouse and at least one child (left truncation) and that individuals could be right censored. We furthermore adjusted for religion (UPDB only), sibship size, birth cohort, sex, socio-economic status, mother’s age at birth, birth order, birth intervals, and twin birth since these are known to influence human survival^1^. socio-economic status was constructed according to the Integrated Public Use Microdata Series (IPUMS) occupational coding scheme of 1950 (OCC1950)^62^. Importantly, for the sibling contribution to the cumulative percentile analysis (analysis 1), the sibling contribution to the top 10% analyses (analysis 3), and in all mutually exclusive group analyses (analysis 3), we used analytical weights when fitting the Cox models to avoid family size confounding. Adjustment was not necessary for the number of parents because this number is two by definition. However, sibship sizes vary. For example, a hypothetical IP with 4 siblings belonging to percentiles 1, 6, 8 and 30 will contribute with a weight w=3/4 in the first analysis, based on the cumulative percentiles, when considering the top 10 percent. This same IP, when considering the top 5 percent will contribute with less weight, namely w=1/4. In this way, each person contributed the same to the overall analysis across all percentiles. In the second analysis based on mutually exclusive groups, this same hypothetical IP would be assigned to g1, and will contribute to the analysis with a weight w=1/4. In analysis 3, based on the top 10%, the IP will contribute with a weight of w3/4 In this way we avoid a potential advantage to larger families to be represented in more extreme groups.

#### Verification of the results in a subsequent generation (analysis 4)

To verify our results regarding the top 10% parents and siblings (analysis 3) in a subsequent generation (children, F3), we investigated whether children of top 10% IPs had a survival advantage compared to children of non-longevous IPs and whether this effect is stronger if the spouse of the IP also belonged to the top 10%. We further investigated familial clustering of longevity by studying the number of top 10% aunts and uncles of the children of IPs. A Cox-type random effect model was used:

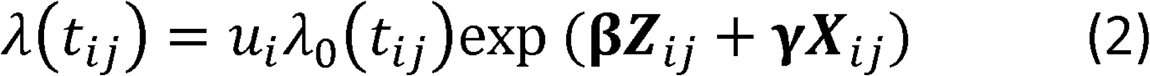

where *t_ij_* is the age at death or the age at last follow-up for child *j* in family *i*, λ_0_(*t_ij_*) refers to the baseline hazard, which is left unspecified, **β** is a vector of regression coefficients for the main effects of interest (***Z***) which correspond to: (1) having a parent top 10% survivor in a first analysis and (2) the effect of the number of uncles/aunts (F2) top 10% in a second analysis. *u* > 0 refers to an unobserved random effect (frailty) shared by F3 children of a given IP. This unobserved heterogeneity shared within sibships was assumed to follow a log-normal distribution. **γ** contains the effect of person-specific covariates ***X***, similar to those included in the previous analyses.

#### Survival of the spouses by the longevity of the index persons (analysis 5)

To investigate the survival of spouses, we applied a group approach, similar to that used above, and analyzed the groups with Cox regression. We grouped the spouses by the survival of the IPs creating 6 different groups g (g = 1,2,3, …, 6): group 1 = [≥0^th^ & ≤1^th^ percentile], group 2 = [≥1^th^ & ≤5^th^ percentile], group 3 = [≥5^th^ & ≤10^th^ percentile], group 4 = [≥10^th^ & ≤15^th^ percentile], group 5 =[≥15^th^ & ≤20^th^ percentile], group 6 = [≥20^th^ & ≤100^th^ percentile]. We compared the groups in two steps: (1) group 6 was the reference category and (2) comparing all groups with each other (post-hoc), applying a Bonferroni correction for multiple testing.

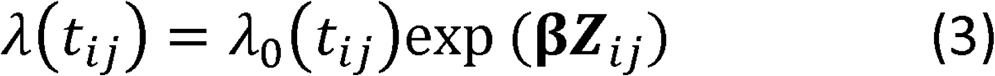

where *t_ij_* is the age at death or the age at last follow-up for spouse *j* in family *i*. λ_0_(*t_ij_*) refers to the baseline hazard, which is left unspecified in a Cox-type model. **β** is the regression coefficient referring to the main effects of interest (***Z***), which are the spouses who were divided into mutually exclusive groups by their parental mortality.

## Acknowledgment

This research was supported by Netherlands Organization for Scientific Research [360-53-180], the Professor van Winterfonds, and the Leiden University Fund [6223/07-06-2016]. I thank Professor Ken R smith for hosting my stay in Utah (US) to work with the Utah Population Database. I also thank Dr. Joris Deelen and Dr. Thies Gehrmann for providing their input to this work as independent readers.

## Author contributions

Niels van den Berg is the study investigator and was responsible for initiating the study, data management, data analyses, writing the first draft of the manuscript and finalizing it, and obtaining funding to visit Ken R. Smith in Utah. Angelique Janssens and P. Eline Slagboom are the study principal investigators who conceived and obtained funding for the project, which this study is a part of. Ken R. Smith is the head of the UPDB and he hosted the stay of Niels van den Berg in Utah, provided access to the UPDB, and supervised all that concerned the UPDB within this study. P. Eline Slagboom and Marian Beekman provided overall project coordination and supervision. Kees Mandemakers is the head of LINKS and provided access and support to the LINKS data. Rick Mourits provided access to documentation and conversion tables of Socio Economic Status coding schemes between the two databases. Ingrid van Dijk assisted with the initial family re-construction process and assisted in working with the LINKS data.

## Data availability

The LINKS data is available upon request to Kees Mandemakers who is the director of the LINKS data at the International Institute of Social History, Cruquiusweg 31, 1019 AT Amsterdam, the Netherlands. The UPDB is also available upon request to Ken R Smith who is the director of the UPDB at the University of Utah, 225 S. 1400 E. Rm 228 Salt Lake City, United States.

